# Context effects on repair of 5’-overhang DSB induced by Cas12a in Arabidopsis

**DOI:** 10.1101/2024.08.05.606594

**Authors:** Sébastien Lageix, Miguel Hernandez Sanchez-Rebato, Maria E. Gallego, Jérémy Verbeke, Yannick Bidet, Sandrine Viala, Charles I. White

## Abstract

Sequence-specific endonucleases have been key to the study of the mechanisms and control of DNA double-strand break (DSB) repair and recombination and the availability of CRISPR-Cas nucleases over the last decade has driven rapid progress in understanding and application of targeted recombination in many organisms, including plants. We present here an analysis of recombination at targeted chromosomal 5’overhang DSB generated by the FnCas12a endonuclease in the plant, *Arabidopsis thaliana*. The much-studied Cas9 nuclease cleaves DNA to generate blunt-ended, double-strand breaks (DSB), but relatively less is known about the repair of other types of breaks, such as those with 5’-overhanging ends. Sequencing the repaired breaks clearly shows that the majority of repaired DSB carry small deletions and are thus repaired locally by End-Joining recombination, confirmed by Nanopore sequencing of larger amplicons. Paired DSB generate deletions at one or both cut-sites, as well as deletions and reinsertions of the deleted segment between the two cuts, visible as inversions. While differences are seen in the details, overall the deletion patterns are similar between repair at single-cut and double-cut events, notwithstanding the fact that only the former involve cohesive DNA overhangs. A strikingly different repair pattern is however observed at breaks flanked by direct repeats. This change in sequence context results in the presence of an alternative class of repair events, corresponding to highly efficient repair by Single-strand Annealing recombination.

## Introduction

DNA double-strand breaks (DSB) are highly toxic lesions, causing both local damage and separating the chromatid arm distal to the break from its centromere. This acentric chromatid can be subsequently lost, leading to loss of many genes. Alternatively either of the broken DNA ends can recombine with other sequences in the genome, to generate major structural genome rearrangements (dicentric/acentric chromosomes, deletions, translocations, inversions). A number of recombination pathways repair DSB very efficiently in living cells and the specific outcome of a given repair event is determined by the recombination mechanism involved in its repair. Thus, in somatic cells of animals and plants, chromosome breaks are predominantly repaired by DNA end-joining recombination (EJ) which, while it frequently results in mutations at the repaired break-site, avoids structural genome damage by restricting repair to sequences immediately flanking the break. Readers are referred to excellent reviews on this subject (Chang et al., 2017; Kamoen et al., 2024; Knoll et al., 2014; Scully et al., 2019; Wright et al., 2018).

The possibility to specifically induce *in vivo* breakage of a chromosomal target sequence has been key to understanding of DSB repair and its consequences and applications. Starting with the use of natural and “designer” endonucleases to cleave specific chromosomal targets, this work has been tremendously accelerated in recent years by the availability of CRISPR/Cas endonucleases. Following on from earlier work using transgenic tester loci, Zinc-finger and TALE nucleases, the application of CRISPR/Cas tools to plant models over the last decade has enabled exciting new experimental approaches, and resulted in major advances in the understanding and application of DNA repair and recombination in model and crop plants (Atia et al., 2024; Fauser et al., 2014; Gao, 2021; Jiang et al., 2013; Kamoen et al., 2024; Li et al., 2013; Nekrasov et al., 2013; Přibylová and Fischer, 2024; Schiml et al., 2016; Shan et al., 2013; Vu et al., 2017; Wolter et al., 2021; Xue and Greene, 2021).

The majority of studies DSB-induced DNA repair/recombination in plants in recent years have involved the Cas9 endonuclease and its derivatives. Sequentially cleaving the two strands of the target DNA, Cas9 generates a blunt-ended or 1-base overhang DSB (Jinek et al., 2012; Kumar et al., 2023; Longo et al., 2024; Swarts and Jinek, 2018; Weiss et al., 2024). This means that most studies of Cas9-induced breaks involve the study of blunt-ended DSB, without single-stranded (ssDNA) overhangs. The mutant Cas9-nickase protein, which cleaves only one strand, is an exception to this and has notably been used in studies of recombination at paired single-strand nicks (eg. (Bothmer et al., 2017; Fauser et al., 2014; Schiml et al., 2016, 2014)). In contrast, the Cas12a (Cpf1) endonuclease produces a staggered break, with 4-5 base 5’ ssDNA overhangs at the ends (Swarts and Jinek, 2018; Zetsche et al., 2015).

Furthermore, Cas12a has technical advantages which facilitate its use in studies of DSB repair, notably that it depends upon an easily multiplexable, single guide RNA (gRNA) for targeting cleavage (Fonfara et al., 2016; Zetsche et al., 2017).

In this work we present an analysis of the patterns of DSB repair recombination following cleavage at paired genomic target sites by FnCas12a in the plant, *Arabidopsis thaliana.* The majority of repaired DSB are repaired locally by End-Joining recombination. While differences are seen in the details, overall the deletion patterns are not greatly affected by differing local sequence contexts flanking the DSB. The presence of flanking direct repeats sequences does however strongly impact the repair pattern - corresponding to highly efficient repair by Single-strand Annealing recombination.

## Materials and Methods

### Genomic target loci

To investigate endogenous repair profiles of CRISPR/Cas12a induced DSBs, we screened the *A. thaliana* genome sequence and selected pairs of target sites at two distinct genomic loci. The first target locus, located in the middle of the left arm of chromosome 1, corresponds to the gene *LPP2* (At1G15080). gRNAs were designed to induce two DSBs in the first intron and second exon respectively (see Figure 1a). The second target locus, Chr3_92, was identified by scanning the TAIR10 Arabidopsis genome sequence with the Tandem Repeat Finder (trf version 4.09; https://tandem.bu.edu/trf/trf.html) (Benson, 1999). Chr3_92 lies in the repeat-rich, pericentromeric region of Chromosome 3 and consists of two perfect 142 bp tandem direct repeats separated by 742nt. The 142 bp repeats are themselves part of two degenerate 428 bp tandem repeats. Two unique Cas12a targets were chosen in the sequences between the two repeats. The Cas12a target sites in this locus are thus flanked by tandem direct repeat DNA sequences (see Figure 5a).

**Figure 1.**
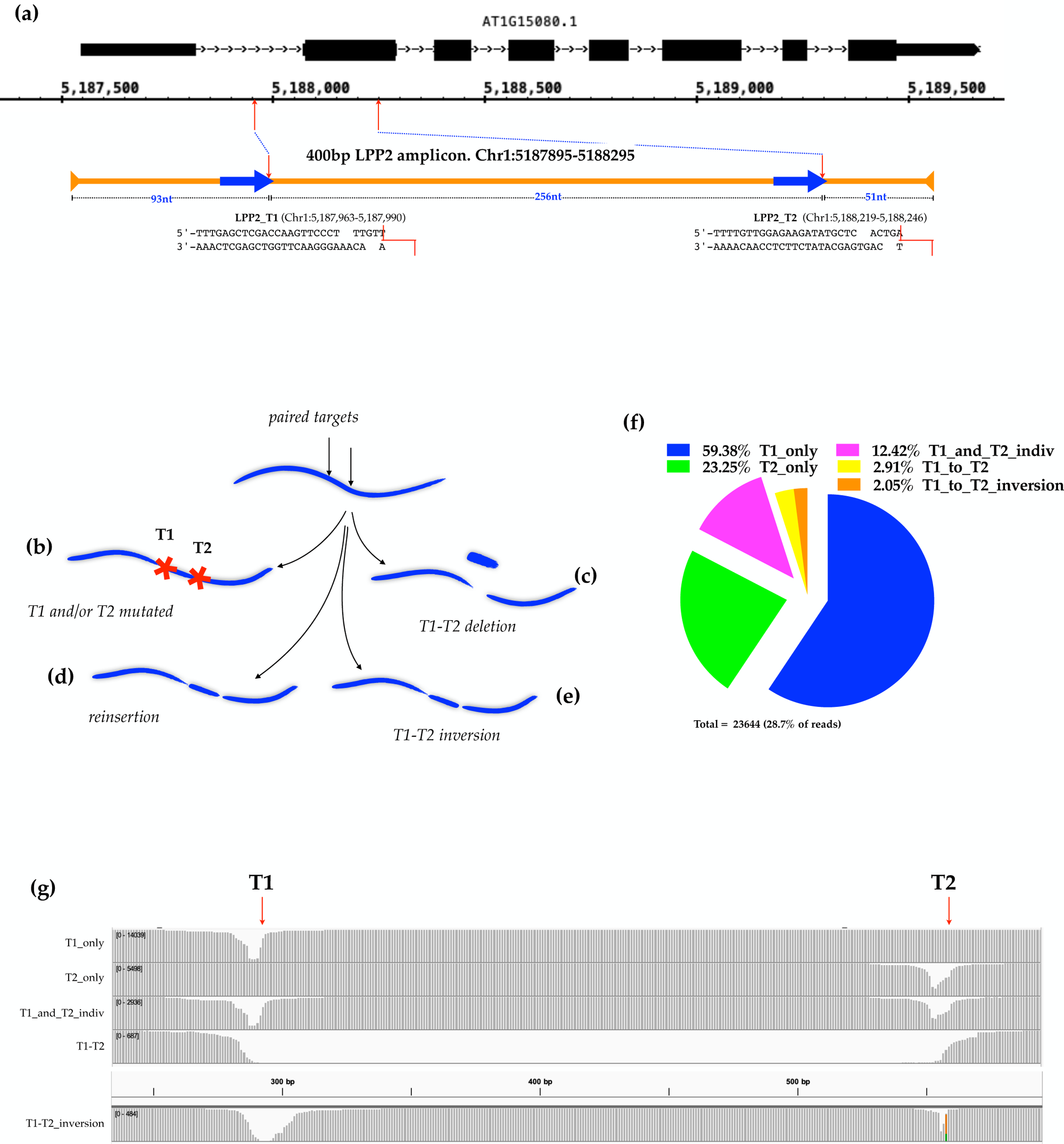
Patterns of repair of paired, 5’-overhang DNA breaks at the LPP2 locus. (a) A schematic showing the two Cas12a target sites of the target LPP2 locus and the 400 bp amplicon used for paired-end sequencing. The sequence, including the (TTT) PAM and the 23nt protospacer, of the targets is shown along with staggered red lines indicating the expected cleavage of the target DNA strands. Cleavage of one or both targets is expected to result in mutation of one or both targets (b). Coincident cleavage at both sites can result in deletion of the DNA fragment between the two targets (c; T1-T2 deletion), possibly accompanied by its reinsertion in either the original (d; not distinguishable from T1 and T2 mutated class (b)), or inverted (e; T1-T2 inversion) orientation. (f) Sorting the paired-end sequences spanning the target locus confirmed the existence and relative proportions of these different classes of event induced in the genome of plants expressing Cas12a. (g) The validity of the sorting is confimed by mapping the sequences of the different repair classes to the Arabidopsis genome, or to an artifical *in silico* inversion locus for the T1-T2_inversion class (bottom).

### T-DNA constructs and plant transformation

Recombinant CRISPR/Cas9 and sgRNA constructs (Table S1) were designed for unique genomic positions and cloned into the expression plasmids kindly provided by Dr Seiichi Toki (Endo et al., 2016). T-DNA transformation of *Arabidopsis thaliana* plants of ecotype Columbia-0 by *Agrobacterium tumefaciens* strain C58C1 was by the floral-dip method (Bechtold, N. et al., 1993; Clough and Bent, 1998).

### Plant selection

In order to obtain transgenic plant lines to address specifically each of the two selected targets, transformed T1 plants for each construct were grown and selected on Murashige and Skoog (MS) medium (4.9 g.l^-1^ Murashige and Skoog medium, 10 g.l^-1^ sucrose, and 8 g.l^-1^agar, pH 5.7) with kanamycin (50 μg.mL^-1^) in a growth chamber with a 16/8-hour light/dark cycle, temperature 23°C and 60% relative humidity. The progeny of individuals T1 were grown for 10 days at 29°C on MS media and gDNA was extracted using Promega’s Wizard Genomic DNA Purification Kit. To assess the cutting efficiency of the transgenic lines, a PCR amplifying the deletion between the two cuts sites for each target loci was performed. The T1 harboring the stronger activity for each construct was selected.

### Sequencing of DNA

The respective target regions from 5 pooled transgenic plants each were amplified by PCR (primer in Table S2) using Platinum SuperFi II DNA Polymerase (Invitrogen) and subjected to NGS. Two different methods were used for NGS applications, the Illumina MiSeq and the Nanopore Technology. The starting material for library preparation was 100 μL of PCR products at 10 ng·μL^−1^. The Illumina MiSeq Nano V2 was used for paired end sequencing of amplicons of a size of 400 bp. For the Nanopore Technology, library preparation was conducted using the SQK-LSK110 ligation sequencing kit following the manufacturer’s protocol. Samples were sequenced using a minION Mk1c device using R9.4.1 (FLO-MIN106D) flow cells.

### Informatics analyses

In-house Python and R scripts using publicly available tools were written to analyse the DNA sequencing data produced in this study (https://github.com/chrlesw/dsbrepair.git). Briefly, the scripts analyse NGS DNA sequence outputs from both Illumina paired-end sequencing or Oxford Nanopore sequencing. The Nanopore sequences require no pre-processing, but prior to running the analyses, the paired-end Illumina sequences are merged into individual sequences using the BBMerge tool of the BBTools package (BBMap – Bushnell B. – sourceforge.net/projects/bbmap/ Brian Bushnell (Bushnell et al., 2017)). These merged sequence files are the input for two main Python scripts, which carries out the analysis following two approaches:

### 1) Text-recognition approach: _dsb_sortAnal.py

Based on text searches and precise sequence lengths, this approach is best applied to the Illumina sequences due to the lower sequencing error rate compared to Nanopore sequences. The merged Illumina sequences are individually scanned for the presence/absence of the Cas12a target sequence(s) using Python text-search tools. The presence of inversions is identified by the presence of a reverse-complementary sequence taken from between the target sites. These, combined with their lengths, are used to sort and enumerate the merged sequences of different deletion classes: single site deletions, individual two-site deletions, deletions spanning the two cut-sites and inversions of the sequences between the two cut-sites.

### 2) Mapping plus variant calling approach: _dsb_varAnal.py

Minimap2 (Li, 2018) is used to align the Nanopore or merged, paired-end Illumina sequences to the reference genome. Secondary or supplementary alignments are removed and the BBMap callvariants.sh script of the BBTools package (BBMap – sourceforge.net/projects/bbmap/ Brian Bushnell (Bushnell et al., 2017)) used to produce lists of deletions, insertions and substitutions. Shell commands then select variants affecting 10 nt windows centered on the cut-sites and sort them into lists of deletions, insertions and substitutions. The potential implication of DNA sequence microhomologies is determined for each deletion junction and plots showing each deletion as an arc linking its start and end points are generated.

## Results

A FnCas12a construct expressing guide RNAs targeting two sites, LPP2_T1 and LPP2_T2, respectively located in the first intron and second exon of the Lipid Phosphate Phosphatase 2 gene (LPP2, AT1G15080). The key characteristic of this gene as a target for this work is that it is non-essential (Katagiri et al., 2005). The two target sites, T1 and T2, are 256 bp apart, oriented in the same direction and both are unique in the *Arabidopsis thaliana* genome (Figure 1a). Stably transformed lines carrying this construct were produced by T-DNA transformation and a line showing high levels of mutation at the target sites was chosen for further work. In parallel a plant line carrying the same FnCas12a T-DNA construct, but lacking the gRNA template was built as a -gRNA control.

Genomic DNA was extracted from 10-day old plantlets, a 400bp region including the target sites was amplified (Figure 1a) and paired-end sequenced with an Illumina Miseq sequencer. The read-pairs were merged with the BBMerge script of the BBMap package to give single sequences and these were analysed by an in-house Python script (see Materials and Methods).

In principle, repair of the paired Cas12a-induced breaks can result in a number of outcomes: simple religation (not detectable) or mutation of either or both T1 or T2 (Fig. 1b), deletion from T1 to T2 (Fig. 1c), T1+T2 cleavage followed by re-insertion of the cut-out fragment in either the original (Fig. 1d), or inverted (Fig. 1e) orientation.

A python script was written to quantify and sort the sequence reads into these classes. Based on the sequence length plus the presence/absence of the unmodified target sequences, this script gives a direct read-out of the events in each sequence. In the Cas12a+gRNA plants, 23644 sequences with modified T1 and/or T2 were found in the 82431 sequence reads (28.7%), while 11 were found in the 52806 reads (0.02%) from the Cas12a-gRNA controls. The proportions of the different classes of events found in the +gRNA sequences are shown in Figure 1f. Thus, of the 28.7% of sequences which carry a deletion in T1 and/or T2, individual T1 deletions (59.38%) are 2.6-fold more frequent than individual T2 deletions (23.25%), presumably due to more efficient cleavage at the T1 target. 12.42% of the sequences with T1 and/T2 deletions have individual mutations at both T1 and T2, and 2.91% deletions extending from T1-T2, with slightly lesser number (2.05%) having T1-T2 deletions associated with reinsertion of the intervening sequence in inverted orientation. It is important to keep in mind that the frequencies of the latter two classes (the T1-T2 events) will depend upon the likelihood of a given chromatid being cleaved at both sites co-incidentally, and thus the relative and absolute frequencies of cleavage at T1 and T2. The relative frequencies of these paired-cut events would thus be expected to differ in different transformants, target pairs or at different loci. Aligning the sorted sequence files of each class of events to the Arabidopsis genome (or an in-silico T1-T2 inversion locus for the inversion sequences) confirms the efficacy of this approach and gives a first view of the nature of the deletions in each class (Fig. 1g).

To take this analysis further, a second script was written to align the merged sequence-pairs to the Arabidopsis genome and call variants with the callvariants.sh script of the BBMap package (see Materials and Methods). Of 83387 sequences mapped to the LPP2 locus from the Cas12a+gRNA line, deletions affecting one or both target sites represent 32.7%, insertions 1.0% and substitutions 4.5% (Figure 2a). As expected, very few deletions (0.02%) or insertions (0.02%) were found in the 52806 sequences from the Cas12a-gRNA control, however numbers of substitutions affecting the target windows were similar in the sequences from the control and Cas12a+gRNA plants (3.3% versus 4.5%). Expressing these numbers as percentages of detected variants (Figure 2b) gives 85.5% deletions, 2.7% insertions and 11.8% substitutions in the +gRNA plantlets, and 0.58% deletions, 0.46% insertions and 98.96% substitutions in the - gRNA controls. The majority of substitutions are thus not Cas12a targeted and presumably arise from sequencing errors, which occur at comparable frequencies in this material.

**Figure 2.**
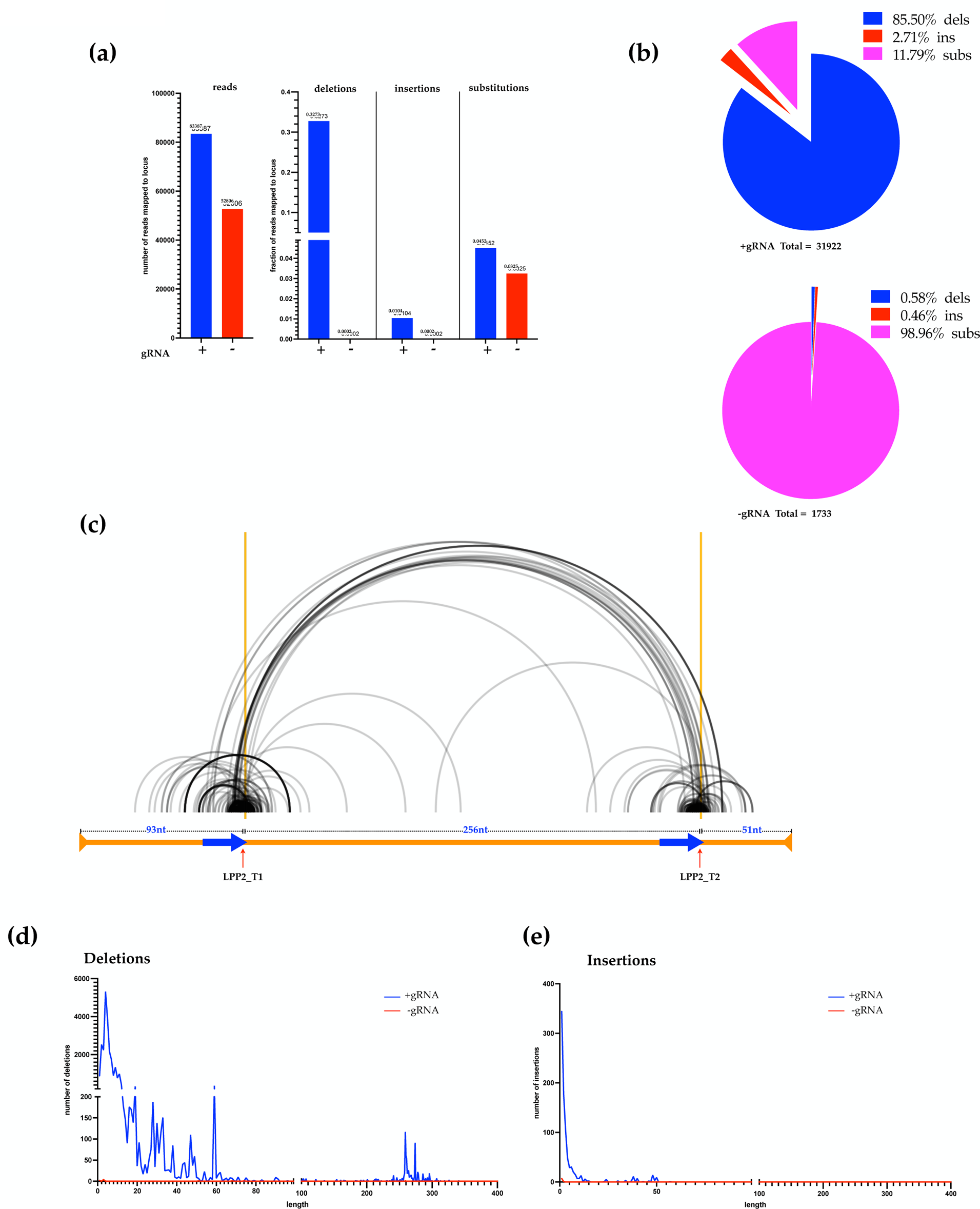
Deletions predominate in the repair of 5’-overhang DNA breaks at the LPP2 locus. (a, b) Analysis of variants affecting 10 bp windows centered on the cut-sites confirms the predominance of deletions in the repair products of Cas12a-induced breaks at the LPP2 locus. DNA from the -gRNA control plants contains almost exclusively base-substitutions, presumably resulting from sequencing errors. (c) Plotting 1000 randomly-chosen deletions as arcs (arcs from start to end of deletion) across the target locus (X-axis, horizontal orange bar), shows that the deletions are almost exclusively focussed locally at the cut-sites or span between them. This is confirmed by plotting the distributions of the numbers of deletions (d) and insertions (e) by their lengths (blue curves are +gRNA and red are -gRNA controls). Vertical orange lines in (c) mark the target-sites.

The great majority of mutations induced by Cas12a at this locus are deletions and are mostly short deletions. A clear, visual confirmation of this is shown in Figure 2c, which shows 1000 randomly-chosen deletions, each plotted as an arc joining its start- and end-points. Both the clusters of small deletions at each cut-site and the larger deletions involving both cut-sites are clearly visible (black arcs). It is particularly striking to note the relatively small number of other deletions extending into the flanking chromosomal DNA (Figure 2c). Frequency distributions of deletion lengths confirm this observation, showing clearly that, with the exception of the T1-T2 deletions from 256-300bp, the great majority are less than 20bp, with only rare events being longer than 60bp (Figure 2d). A similar analysis shows that practically the totality of insertions are 5bp or less, with a small minority extending to 10bp (Figure 2e). The FnCas12a endonuclease cleaves double-stranded DNA (dsDNA) to leave 4-5nt, 5’ single-strand (ssDNA) overhangs (Zetsche et al., 2015) and we expected to find a major class of 4-5nt deletions corresponding to loss of the ssDNA overhangs at the cut sites, followed by religation. Frequency distributions of 4 and 5nt deletion start- and end-sites confirm the presence of this class of “blunting/ligation” NHEJ events (Figure 3a). This is particularly clearly visible at the T1 site, where practically the totality of 4- and 5-nt deletions are at the expected position. It is also seen at the T2 site, however deletions starting 2 and 3 nt upstream of the expected peak are also clearly visible. Should these upstream deletion start sites result from extended erosion of the upstream sequence prior to religation, this would result in deletions longer than the 4-5bp and their exclusion from this analysis. We can thus confidently class these events as resulting from other cleavage sites upstream of the “primary” cut site.

**Figure 3.**
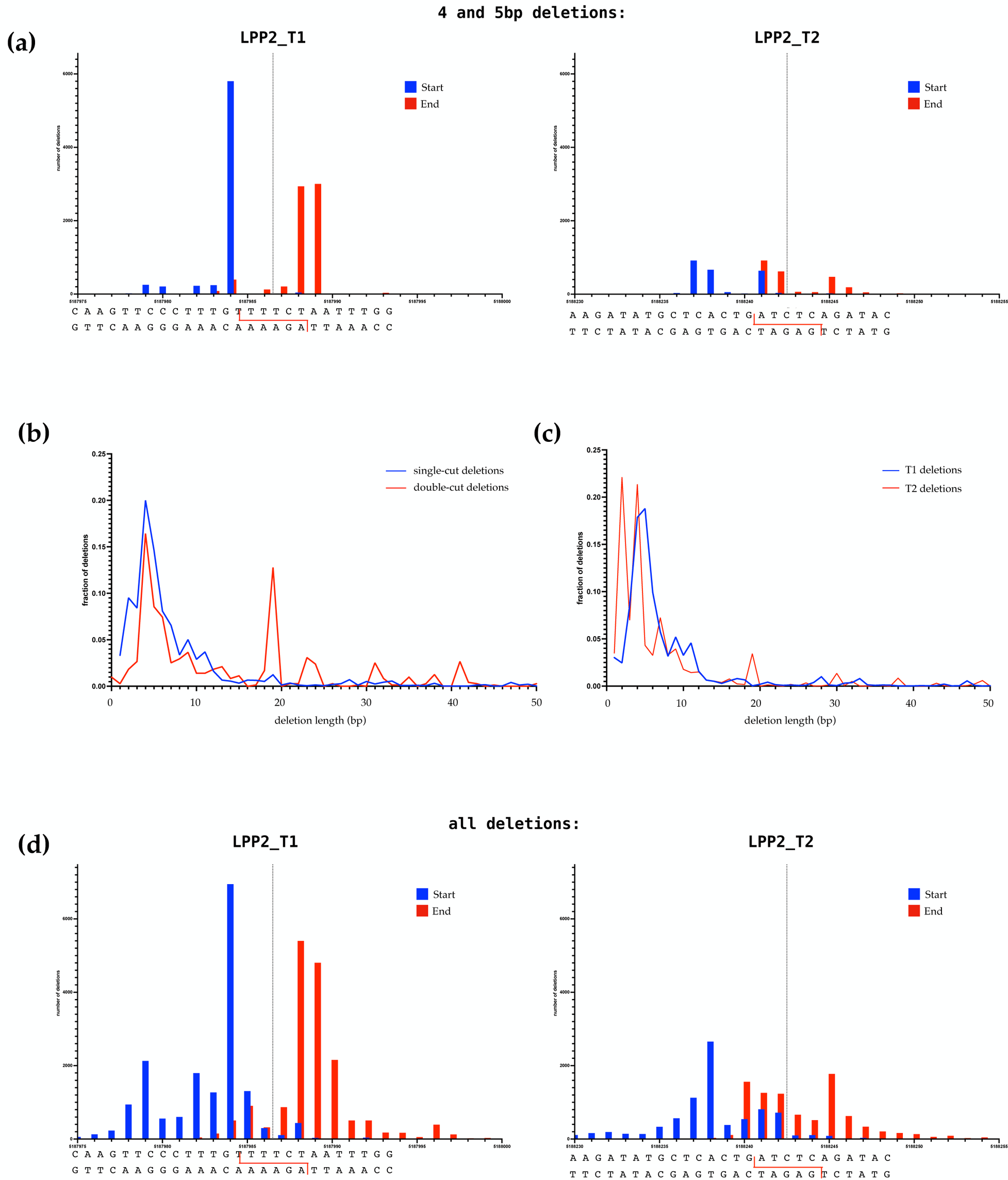
Patterns of deletions at the LPP2_T1 and LPP2_T2 targets. (a) Frequency distributions of start-sites (blue) and end-sites (red) of 4 and 5nt deletion affecting the 10 bp windows centered on the cut-sites. The red bars have been offset slightly to the right for clarity. The target locus sequence with the cut-sites marked by staggered red lines, is shown below the graph. Vertical dotted lines show the centers of the cut-sites to facilitate lecture of the graphs. The majority of these deletions begin at the base pair immediately upstream of the 5’ ssDNA overhang produced by Cas12a cleavage of the LPP2_T1 target, and the end points are found 4 and 5 bases downstream. The LPP2_T2 target has 3 preferred start-points for 4 and 5nt deletions, including deletions starting 2 and 3 nt upstream of the expected cut. (b) Frequency distributions of lengths of single-cut and double-cut (transposed 256nt leftwards) confirm that the deletion patterns of the majority, small local deletions aren’t affected by the presence of cohesive (1-cut) versus non-cohesive (2-cut) 5’ ssDNA overhanging ends. Taken individually, frequency distributions of deletion lengths (c) and deletion start- and end-points (d; same presentation as in (a)) at the two cut-sites are dominated by short local events, notwithstanding the presence of differences in the details of the distributions at the two sites.

Each Cas12a cut produces complementary and potentially religatable overhanging 5’ ssDNA ends, however this is not so for the double-cut events (deletions ≥256bp), which involve joining of DNA ends from cuts in two different sequences. This distinction does not appear to significantly affect the loss of sequence in the deletions, as can be seen by overlaying the distributions of the 1-cut and the 2-cut events (by subtracting 256 from the lengths of the 2-cut deletions) (Figure 3b). To facilitate this comparison, the frequency distributions in Figure 3b are presented as fractions of total deletions. Plotting the distributions of T1 and T2 deletions separately confirms that, while overall very similar, unsurprisingly there are differences in the details of the patterns of deletion lengths (Figure 3c) and the choice of start and end-points (Figure 3d) at the two target sites. It appears reasonable to assume that this is at the origin of the minor differences between the patterns of local sequence loss in the single- and double-cut EJ events (Figure 3b). Furthermore, given that the targets both lie within the LPP2 gene, separated by only 256bp, these specificities are presumably due to local DNA sequence context. Finally, the maximum number of nt of microhomology from 0 to 9 nt, potentially involved in each deletion was calculated (see Figure S1a). Mean microhomology values for all deletions starting at each coordinate were calculated across the LPP2-T1 and LPP2_T2 target loci (Figure S1b and c). The number of deletions starting at each coodinate was also plotted to put these values into context, confirming the absence of correlation between microhomology and deletion frequency.

Due to restrictions on amplicon length for Illumina sequencing, the cut-sites analysed above are 93 and 52bp from the ends of the PCR-amplified fragment used for the sequencing (Figure 1a). Loci with deletions extending beyond these ends would not be amplified (nor sequenced) and this could clearly impact our analyses. We thus used Oxford Nanopore sequencing, which doesn’t suffer from these constraints on amplicon sequence length, to test whether the Illumina approach had impacted our analyses and, if so, to build a clear picture of the DSB-induced recombination at this locus.

A 2.8kb *LPP2* amplicon was chosen for Nanopore sequencing (Figure 4a). In this case, the T1 and T2 targets are respectively 1319bp and 1226nt from the amplicon ends. This segment was amplified from 10-day old plantlets expressing the Cas12a+gRNA (targeting LPP2 as above) and from plantlets expressing Cas12a+gRNA targeting another locus on chromosome 3 as Cas12a-gRNA control (Chr3_92, see below). Although the Nanopore sequencing data includes a significant background of small deletions and insertions, these are almost all less than 6bp long (Figure 4b, c), and when excluded from the analysis, the deletion patterns from the Illumina and Nanopore sequencing overlap very closely (Figure S2c).

**Figure 4.**
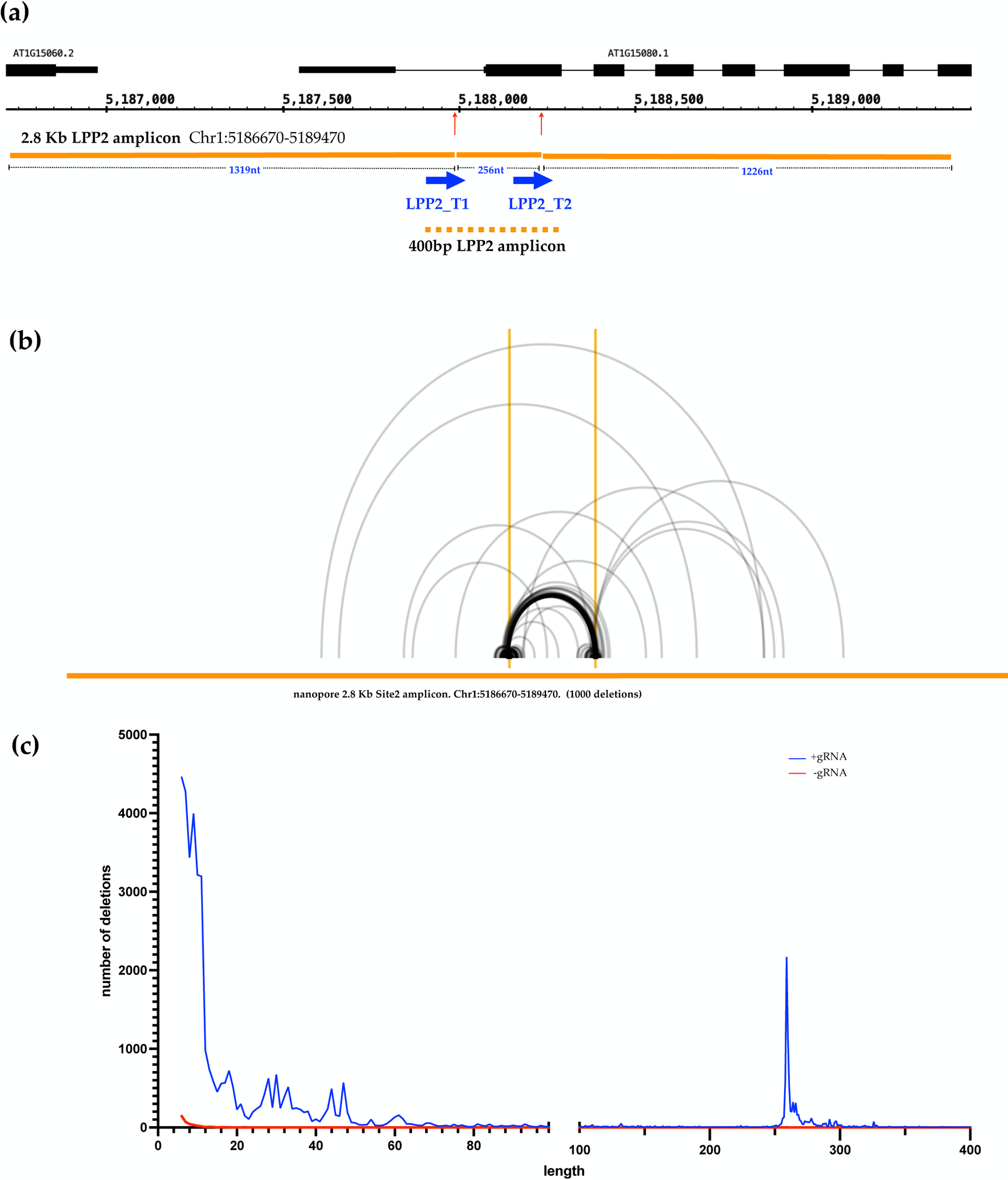
Nanopore sequencing confirms patterns of Cas12a-induced deletions at the LPP2 targets. (a) Schematic showing the two Cas12a target sites of the target LPP2 locus and the 2.8 Kb amplicon used for Nanopore sequencing. (b) Plot showing arcs linking each deletion start- and end-point across the target locus (X-axis, horizontal orange bar). Vertical orange lines mark the target-sites. The arcs are all of the same grey colour, the darker lines reflect overlaying of multiple arcs. (c) Distributions of numbers of deletions by their lengths in Cas12a+gRNA plants (blue curve) and Cas12a-gRNA controls (red curve). Deletions of less than 5 bp were excluded due to the high background of small indel errors in Nanopore sequences.

Of 85439 sequences mapped to the LPP2 locus from the Cas12a+gRNA line, 48587 indels of greater than 5bp affecting one or both target sites were found and of these, 98.7% were deletions and 1.3% insertions (Figure S2a, S2b). As expected, very few indels >5bp (409: 379 deletions and 30 insertions) were found in the 87331 sequences from the Cas12a-gRNA controls (Figure S2b). These numbers from the Nanopore sequencing of the 2.8kb amplicon are thus comparable to those from the Illumina sequencing of the 400bp amplicon.

Although care needs to be taken with the totals due to the exclusion of indels of 5bp or less, Nanopore sequencing of the 2.8kb amplicon does permit detection of longer indels that may have been artefactually excluded from the 400bp amplicon/Illumina approach. Figure 4b presents an Arc-diagram plot of 1000 random deletions of >5bp from the +gRNA Nanopore sequencing and Figure 4c the frequency distribution plot of all detected deletions of >5bp.

Comparing Figure 4b with the equivalent plot from the Illumina sequencing (Figure 2c) shows that, as expected, the longer amplicon Nanopore sequencing does permit detection of deletions extending further into the sequences flanking the cut-sites than those detected with the 400bp amplicon used for the Illumina sequencing. These are, however, clearly only a minority of the deletions and furthermore, are restricted to the central part of the amplified sequence (Figure 4c).

Taken together these results give a clear picture of Cas12a-induced recombination at this locus in Arabidopsis plantlets, showing clearly that repair of the great majority of mutagenic repair of the breaks results in short, local deletions encompassing the breakpoint. To extend these conclusions and test the possible impact of different DNA sequence contexts on the repair mechanisms and outcomes, we applied this approach to a second target locus, in which the Cas12a target sites are flanked by tandem direct repeat DNA sequences. The chosen locus, Chr3_92, lies in the repeat-rich, pericentromeric region of Chromosome 3 (Figure 5a). This locus has two 142nt direct repeats separated by 742nt and two unique Cas12a targets were chosen in the sequences between them. The perfect (no mismatches) 142 nt repeats are themselves embedded in two mildly degenerate (18 mismatches) 428nt direct repeats (Figure 5a).

**Figure 5.**
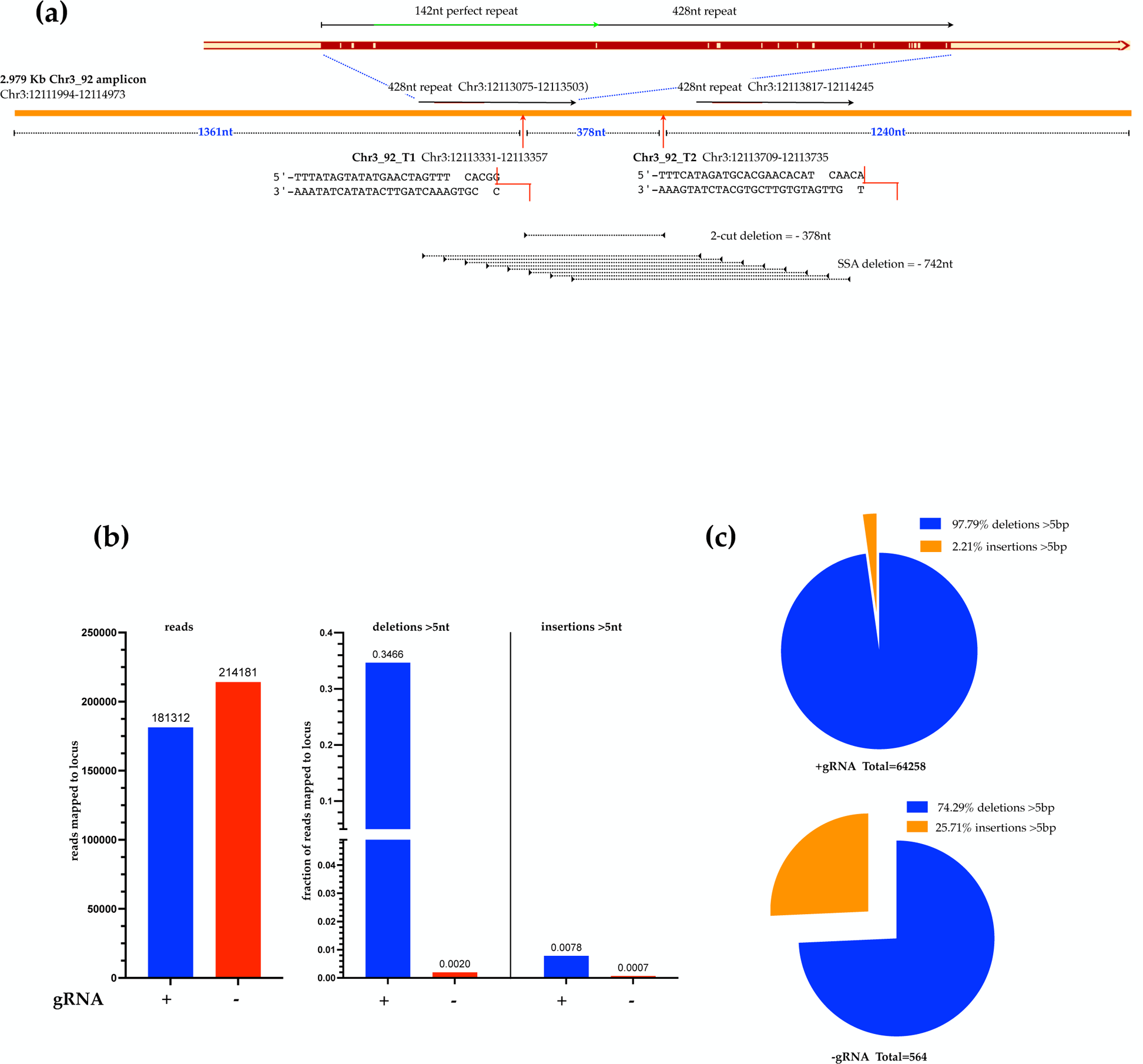
Patterns of Cas12a-induced deletions at the Chr3_92 target locus. (a) Schematic showing the two Cas12a target sites flanked by direct-repeat sequences of the target Chr3_92 locus and the 2.979 Kb amplicon used for Nanopore sequencing. The target sequences, including the (TTT) PAM and the 23nt protospacer, are shown along with staggered red lines indicating the expected cleavage of the target DNA strands. (b) Numbers of sequences and the deletions and insertions expressed as fractions of mapped to the Chr3_92 locus from Cas12a+gRNA (blue) and Cas12a-gRNA plants (red). Indels of less than 5 bp were excluded due to the high background of small indel errors in Nanopore sequences. (c) As seen at the LPP2 locus, the majority of Cas12a-induced indels are deletions (b, c).

Following the same logic as for the analyses at the LPP2 locus, a 2.8Kb amplicon centered on the Chr3_92 locus was amplified from 10 day-old plantlets expressing the Cas12a+gRNA targeting Chr3_92_T1 and Chr3_92_T2 on chromosome 3 and a Cas12a+gRNA targeting another locus as Cas12a-gRNA control (the LPP2 locus on Chromosome 1, see above). The Chr3_92_T1 and Chr3_92_T2 targets, separated by 378nt, are 1361bp and 1240nt from the amplicon ends respectively (Figure 5a). This 2979bp segment (Chr3:12111994-12114973) was amplified by PCR and sequenced with the Oxford Nanopore Minion sequencer (as for the LPP2 locus, above).

Of 181312 sequences mapped to the Chr3_92 locus from the Cas12a+gRNA line, 64258 indels of greater than 5bp affecting one or both target sites were found and of these, 97.79% were deletions and 2.21% insertions (Figure 5b, 5c). As expected, very few indels >5bp (564: 419 deletions and 145 insertions) were found in the 214181 sequences from the Cas12a-gRNA controls (Figure 5b, 5c). These numbers from the Nanopore sequencing of the 2.8kb amplicon are thus comparable to those from the Illumina sequencing of the 400bp amplicon.

Analysis of the Cas12a-induced deletions at the Chr3_92 locus shows a pattern of local deletions, at and between the two cut-sites (Figure 6a), comparable to that observed at the LPP2 locus (Figure 4b). However, in contrast to the LPP2 deletion pattern, a new major deletion class at 742bp represents 39.37% of the 61689 deletions greater than 5bp (Figure 6a, 6b). 742 bp is precisely the spacing between the flanking repeat sequences and these events thus presumably result from SSA recombination. The Arc-plot of the deletions confirms that this is the case: a major class of deletions with start/end points distributed across the repeat sequences, extend in-step from one repeat to the other (red arcs in Figure 6c). The presence of these flanking direct repeat sequences thus generates a new class of recombinants representing almost 40% of the total, that is not detected in their absence. The arc-plot of the LPP2 deletions from Figure 3b is inset in Figure 6c for comparison.

**Figure 6.**
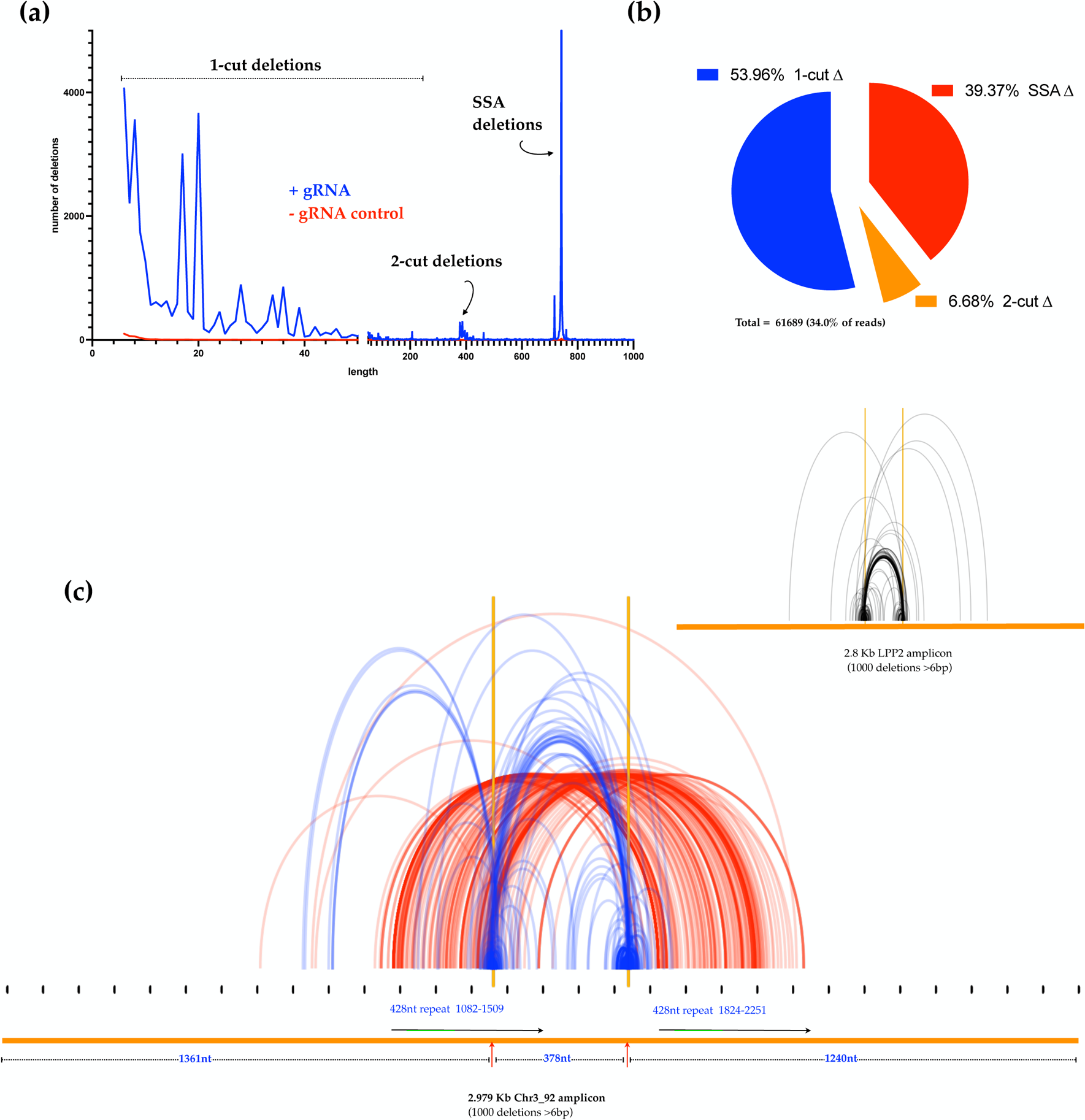
The presence of flanking direct-repeats generates a major new class of Single-Strand Annealing deletions. (a) Distributions of numbers of deletions by their lengths in Cas12a+gRNA plants (blue curve) and Cas12a-gRNA controls (red curve). Deletions of less than 5 bp were excluded due to the high background of small indel errors in Nanopore sequences. A major (39.4%), novel class of 742 bp deletions is created by the presence of the direct repeats flanking the Cas12a target sites. (c) Deletion start-end arc-plots confirm that these 742 bp deletions (red curves) result from in-step recombination between the flanking direct-repeats, as expected from SSA recombination. This is strikingly different from the patters observed at the LPP2 locus in the absence of the flanking repeats (inset in (c), repeated from Figure 4).

The optimum temperature for activity of the FnCas12a endonuclease in plants is 29°C, (Bernabé-Orts et al., 2019; Huang et al., 2021; Lee et al., 2019; Malzahn et al., 2019; Schindele and Puchta, 2019) and the data presented in this article comes from analysis of plants grown at this temperature. While 29°C is a temperature well within the range that Arabidopsis plants will encounter in the environment and the plants grow well at this temperature, the standard laboratory growth temperature for *Arabidopsis thaliana* is 23°. We thus verified that the higher growth temperature doesn’t impact the choice of mechanism and thus outcomes of repair of Cas12a-induced DSB in the plants. Illumina sequencing of the 400bp LPP2 locus amplicon from plantlets grown at 23°C and 29°C shows similar proportions of the different deletion classes (Figure S3a) and the distributions of deletions (Figure S3b) and insertion (Figure S3c) lengths coincide at the two temperatures. The patterns of End-Joining repair of these breaks at 23° and 29° are thus equivalent. We also checked for a possible temperature-specific impact on the SSA recombination pathway by making an analogous comparison between results or repair of DSB targeted to the Chr3_92 locus. As for the results concerning EJ recombination at the LPP2 locus (above), similar proportions of the different deletion classes (Figure S4a) are observed and the distributions of deletion (Figure S4b) and insertion (Figure S4c) lengths overlap at the two temperatures. The patterns of End-Joining and SSA repair of 5’-overhang DSB at these two loci are thus equivalent at these two temperatures.

## Discussion

This study concerns the repair of 5’-ssDNA overhang DSB in *Arabidopsis thaliana*, using targeted *in planta* cleavage by FnCas12a (FnCpf1) (Endo et al., 2016). In addition to producing 5’-ssDNA overhang DSB, Cas12a has high specificity for its target and the very useful capability to process precursor RNAs into crRNAs (Fonfara et al., 2016; Makarova et al., 2015; Shmakov et al., 2015; Strohkendl et al., 2018; Swarts and Jinek, 2018; Zetsche et al., 2015). The presence of 5’ overhangs at Cas12a-induced DSB is of particular interest for study of DNA break repair and recombination. Like blunt-ended and 3’-ssDNA overhang DNA breaks, 5’-ssDNA overhang DSB can be repaired by simple religation. Such events are however very difficult to detect and analyse, as well as being subject to further cleavage/repair.

Degradation of the 4-5nt 5’ overhangs or filling-in of the recessed 3’-ended strands left by Cas12a cleavage to generate blunt ends for EJ ligation would result in 4-5 bp deletions or insertions respectively. Further resection of the 5’-ended strand would be needed to produce the 3’ overhangs needed for microhomology-dependent alternative end-joining (aEJ or MMEJ) or homology-dependent recombination (HR) and single strand annealing (SSA).

As well as being less studied, notably in plants, these breaks are thus of particular interest in terms of recombination. We chose to work with pairs of nearby Cas12a cuts in order to characterise any specificities of repair of the (cohesive-ended) single Cas12a cuts, versus the (non-cohesive-ended) twin-cuts and to check for more complex events involving re-insertion of a cut-out chromosome fragment. We note also that our PCR-amplicon based analysis is necessarily restricted to “local” DSB repair outcomes. While we specifically include analysis of twin-break events and include long-read sequencing to take this into account, larger scale rearrangements (chromosome arm loss, translocations…) are not directly analysed in the work reported here. In this context, we direct readers to the recent study and discussion of Cas12a-induced mutations in the ascomycete fungus *Magnaporthe oryzae* (Huang et al., 2022). Finally, we note that while Cas12a enzymes are strongly impacted by protospacer/target mismatches, they cut the target DNA at the PAM-distal end of the target sequence (see (Strohkendl et al., 2018; Swarts and Jinek, 2018)). Thus, recognition and re-cleavage of repaired targets with small sequence changes (substitutions, small indels) will potentially be more frequent than would be the case with PAM-proximal cleavage by enzymes such as Cas9 (Swarts and Jinek, 2018). While this effet could be expected to lead to longer deletions, our results however show a marked predominance of very short deletions, suggesting that the impact of this in our plants is less than might have been expected, perhaps due to the very efficient cutting.

Five principle conclusions emerge from the work presented here:

1. The great majority of mutations left by the repair of these 5’-overhang DSB are deletions, with a clear peak of deletion lengths at 4bp. The frequencies then fall off rapidly with increasing deletion length. Strikingly, repair resulting in insertions (also very short) is > 30-fold less frequent than that producing deletions. Examination of the patterns of deletion lengths shows them to be very similar at the different cut-sites, with however some differences seen in the details (see below). Plotting mean microhomology length against deletion start point across the target sites shows no clear evidence for a relation between the probability of a given base pair serving as the start-point of a deletion and the mean number of nt of microhomology potentially involved in the deletions starting at that position in the sequence. While the existence of such microhomologies in the sequence in no way proves that they were actually involved in the recombination event under study, this result does indicate that the use of longer microhomolgies is not a primary determinant in the choice of the start and end-points of these very short deletions resulting from repair of 5’-overhang DSB in Arabidopsis.
2. The proximity of the FnCas12a cut-sites to the ends of the 400bp LPP2 amplicon used for Illumina sequencing could clearly impact the patterns of deletions found in this work, excluding repair events involving longer deletions. We thus amplified a 2.8 Kb fragment covering the same locus, for sequencing with the Oxford Nanopore Minion sequencer. The two FnCas12a cut-sites are 1319 and 1226 bp from the ends of this 2.8 Kb amplicon, considerably more distant than the 93 and 51bp in the 400 bp amplicon. Strikingly, while the results of sequencing the longer amplicon did permit identification of longer deletions, these were increasingly rare with increasing length and the overall pattern of repair wasn’t changed relative to that from the 400 bp amplicon sequencing.
3. The use of nearby pairs of Cas12a targets results in a spectrum of modifications at the repaired cuts: individual, local deletions at one or both targets, and deletion and inversions of the DNA sequences between the two targets (Figure 1). While the relative proportions of “one-cut” versus “two-cut” events will be dependent upon the relative and absolute cutting efficiencies at the two sites, examination of the “excision” events involving coincident cutting of both target sites does yield some interesting conclusions. These include the 2.91% of sequences with deletions spanning the two cut sites (T1-T2 deletions) and the events involving reinsertion of the excised T1-T2 fragment. Depending on the orientation of the reinsertion of the excised fragment, the “excision-reinsertion” events would include both the 2.05% of sequences with T1-T2 inversions and an equivalent number classed within the 12.42% of sequences with local deletions at both T1 and T2. Thus the excision-reinsertion class would represent ∼4% of sequences. That this value is comparable in magnitude to the number of T1 to T2 deletions (excision without reinsertion (∼3%) suggests that loss or reinsertion of the excised fragmant in two-site cleavage events are equally likely.
4. The use of paired target sites permitted analysis of patterns of deletions at junctions resulting from joining DNA ends with complementary 5’-overhangs (1-cut deletions) and at those involving non-complementary 5’-overhangs (two-cut deletions). The two cut-sites are separated by 256 bp and subtracting this from the lengths of the T1-T2 deletions permits overlaying the pattern of the T1-T2 deletions with the 1-cut events. While some differences are made clearly visible by this approach, the predominance of 4 bp deletions, followed by a rapid fall-off with increasing deletion-length is strikingly similar. Thus, the presence/absence of cohesive 5’ overhangs does not affect the deletion pattern.
5. The pattern of EJ repair of the 5’-overhang DSB thus results overwhelmingly in the production of very short deletions, with the precise details of the deletion end-points being impacted by local sequence context. This is the case for the two target sites in the chromosome-arm, expressed LPP2 gene and the two targets in the pericentromeric, heterochromatic Chr3_92 locus. The Chr3_92 cut-sites are flanked by tandem direct repeat sequences and this leads to the presence of a new class of deletion events resulting from Single-Strand Annealing (SSA) recombination. It is particularly striking that these SSA events are a major repair class, greatly outnumbering the 2-cut deletions and approaching in frequency the EJ repair events (39% versus 61% of deletions respectively). It is possible that these events result from microhomology-mediate end-joining of a sequence close to the cut site with a target in the flanking repeat. That this is not so is seen clearly in the fact that, rather than being concentrated close to the cut-sites, the start and end-points of the SSA deletions are distributed throughout the (degenerate) repeat sequences. The presence of these flanking direct repeats thus leads to a striking change in the pattern of repair, channeling almost 40% of events into the SSA pathway.

## Acknowledgements

We thank Olivier Mathieu, Yoan Renaud and the members of the recombination group for their help and suggestions.

This work was supported by the Agence National de Recherche (ANR-16-CE91-0010-01 RecInChromatin), the European Marie Sklodowska Curie Actions (EU H2020-MSCA-ITN-2017-765212), the Centre National de Recherche Scientifique (CNRS), the Institut National de la Santé et de la Recherche Médicale (Inserm) and the Université Clermont Auvergne.

## Data Statement

The sequence files are available under BioProject accession number PRJNA1113952 at https://www.ncbi.nlm.nih.gov/sra/PRJNA1113952 (Biosample Accessions: SAMN41471278, SAMN41471279, SAMN41471280, SAMN41471281, SAMN41471282, SAMN41471283, SAMN41488828, SAMN41488829, SAMN41488830).

The Python and R scripts using publicly available tools written to analyse the DNA sequencing data produced in this study can be downloaded from: https://github.com/chrlesw/dsbrepair.git

## Supporting Information

### Supporting Figures and Tables

**Figure S1. Potential implication of microhomologies in deletion formation.**

**Figure S2. Cas12a-induced Indel patterns are confirmed by 2.8 Kb amplicon Nanopore**

**Figure S3. Cas12a-induced Indel patterns at the LPP2 locus from plants grown at 23°C and 29°C.**

**Figure S4. Cas12a-induced Indel patterns at the Chr3_92 locus from plants grown at 23°C and 29°C.**

**Table S1. DNA sequences used for sgRNA constructs.**

**Table S2. List of primers used in this study.**

## Author Contributions

Designed Research and Analysis: SL, MG, YB, MH, CIW; Carried out experiments: SL, MG, MH, SV, JV, CIW; Resources: MG, YB, CIW;

Writing, with contributions from all of the authors: SL, MG, CIW.

## Supporting Figure Legends

**Supporting Figure S1. Potential implication of microhomologies in deletion formation.**

(a) The maximum number of nt of microhomology, from 0 to 9 nt, by comparing the sequences of the upstream DSB-end with the downstream end of the deletion, and the downstream DSB-end with the upstream end of the deletion (black bars connected by dotted lines). The lengths of such microhomologies, potentially involved in the deletion, were determined for each deletion and the mean microhomology values calculated for each deletion start-point across the target sequence. These are plotted as vertical blue bars for the sequences upstream of the LPP2_T1 (b) and LPP2_T2 (c) targets. The number of deletions starting at each start-point are overlaid (red curve) and the cut-site (vertical dotted lines) are included for reference.

**Supporting Figure S2. Cas12a-induced Indel patterns are confirmed by 2.8 Kb amplicon Nanopore sequencing.**

(a) Numbers of sequences and the deletions and insertions expressed as fractions of mapped to the LPP2 locus from Cas12a+gRNA (blue) and Cas12a-gRNA plants (red). Indels of less than 5 bp were excluded due to the high background of small indel errors in Nanopore sequences. As seen from the Illumina sequencing of the 400 bp amplicon (Fig. 2), the majority of Cas12a-induced indels are deletions (a, b). Deletion-length frequency distributions from the 400 bp paired-end Illumina (blue curve) and 2.8 Kb Nanopore (red curve) sequencing data (c).

**Supporting Figure S3. Cas12a-induced Indel patterns at the LPP2 locus from plants grown at 23°C and 29°C.**

Illumina sequencing of the 400bp LPP2 locus amplicon from plantlets grown at 23°C and 29°C shows similar proportions of the different deletion classes (a). This is also seen in the distributions of deletion-lengths (b) and insertion-lengths (c) from plantlets grown at 29° (blue curves) and 23° (red curves).

**Supporting Figure S4. Cas12a-induced Indel patterns at the Chr3_92 locus from plants grown at 23°C and 29°C.**

Nanopore sequencing of the 2.8 Kb Chr3_92 locus amplicon from plantlets grown at 23°C and 29°C shows similar proportions of the different deletion classes (a). This is also seen in the distributions of deletion-lengths (b) and insertion-lengths (c) from plantlets grown at 29° (blue curves) and 23° (red curves).

**Table S1. DNA sequences used for sgRNA constructs.**

**Table S2. List of primers used in this study.**

