## Supplemental Figures and Tables for "Context effects on repair of 5’-overhang DSB induced by Cas12a in Arabidopsis"

### Figure S1

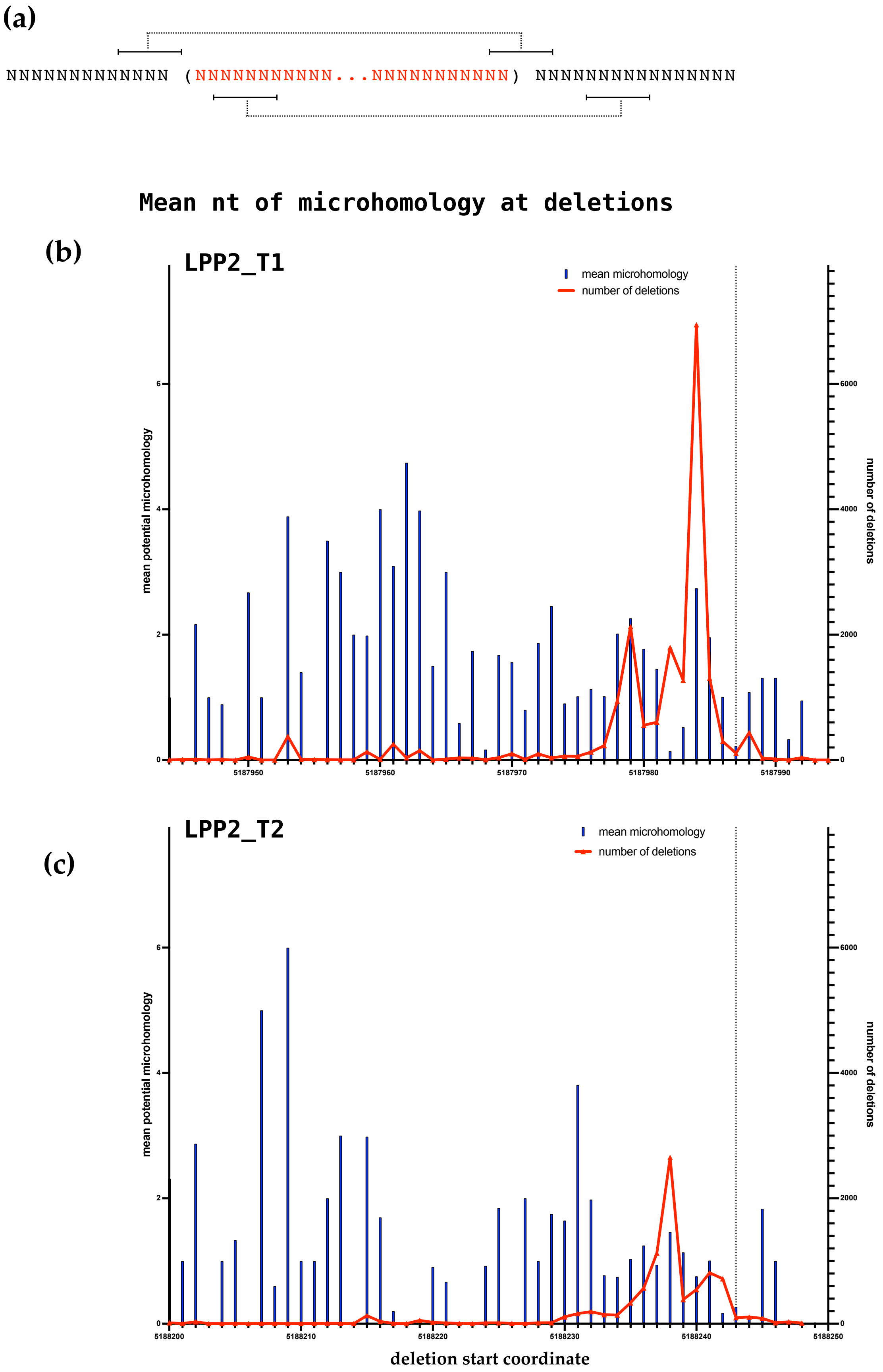
(a)
NNNNNNNNNNNNN (NNNNNNNNNNN...NNNNNNNNNNN) NNNNNNNNNNNNNNNN
Mean nt of microhomology at deletions
(b)
LPP2_T1
mean microhomology
number of deletions
mean potential microhomology
number of deletions
0
2
4
6
0
2000
4000
6000
5187950
5187960
5187970
5187980
5187990
(c)
LPP2_T2
mean microhomology
number of deletions
mean potential microhomology
number of deletions
0
2
4
6
0
2000
4000
6000
5188200
5188210
5188220
5188230
5188240
5188250
deletion start coordinate

#### Figure S2

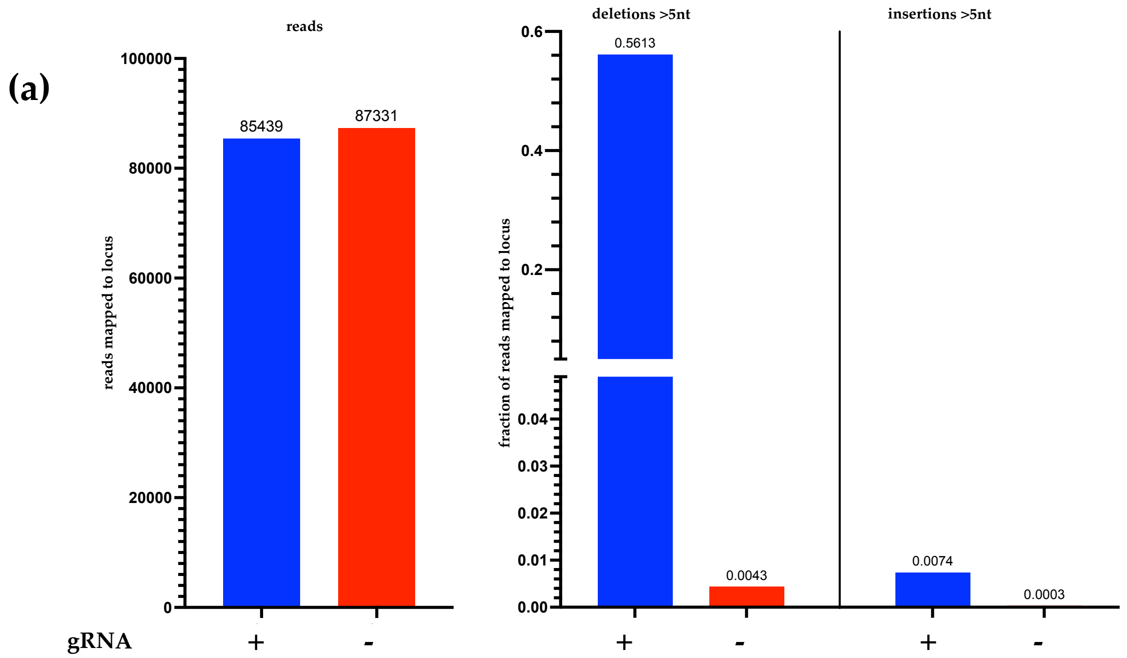
(a)
reads
reads mapped to locus
100000
80000
60000
40000
20000
0
85439
87331
gRNA
+
-
deletions >5nt
insertions >5nt
fraction of reads mapped to locus
0.6
0.4
0.2
0.04
0.03
0.02
0.01
0.00
0.5613
0.0043
0.0074
0.0003
+
-
+
-

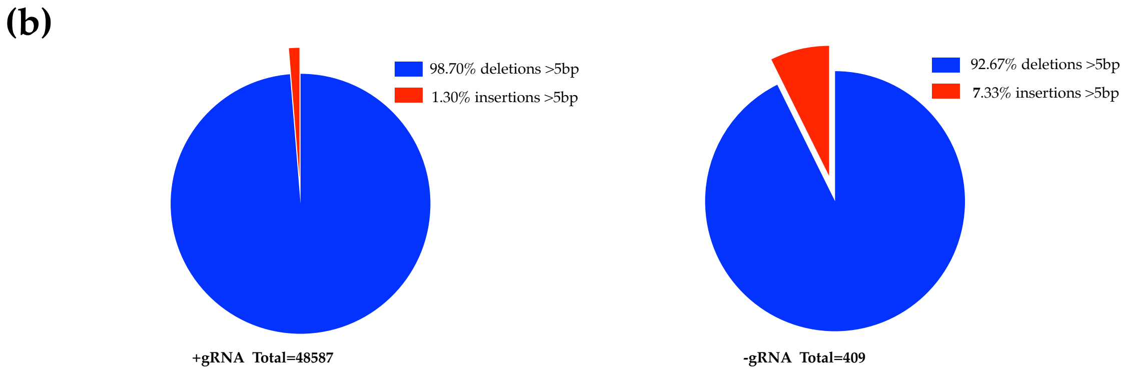
(b)
98.70% deletions >5bp
1.30% insertions >5bp
+gRNA Total=48587
92.67% deletions >5bp
7.33% insertions >5bp
-gRNA Total=409

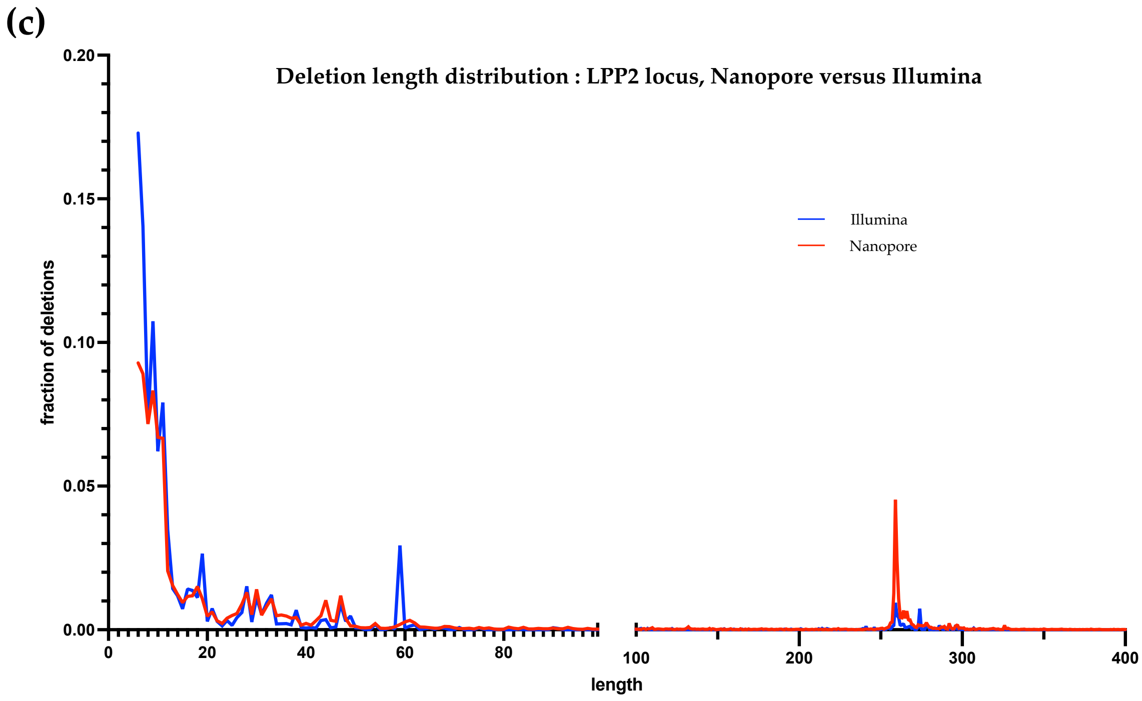
(c)
Deletion length distribution : LPP2 locus, Nanopore versus Illumina
fraction of deletions
0.20
0.15
0.10
0.05
0.00
Illumina
Nanopore
0
20
40
60
80
100
200
300
400
length

**Figure S3**

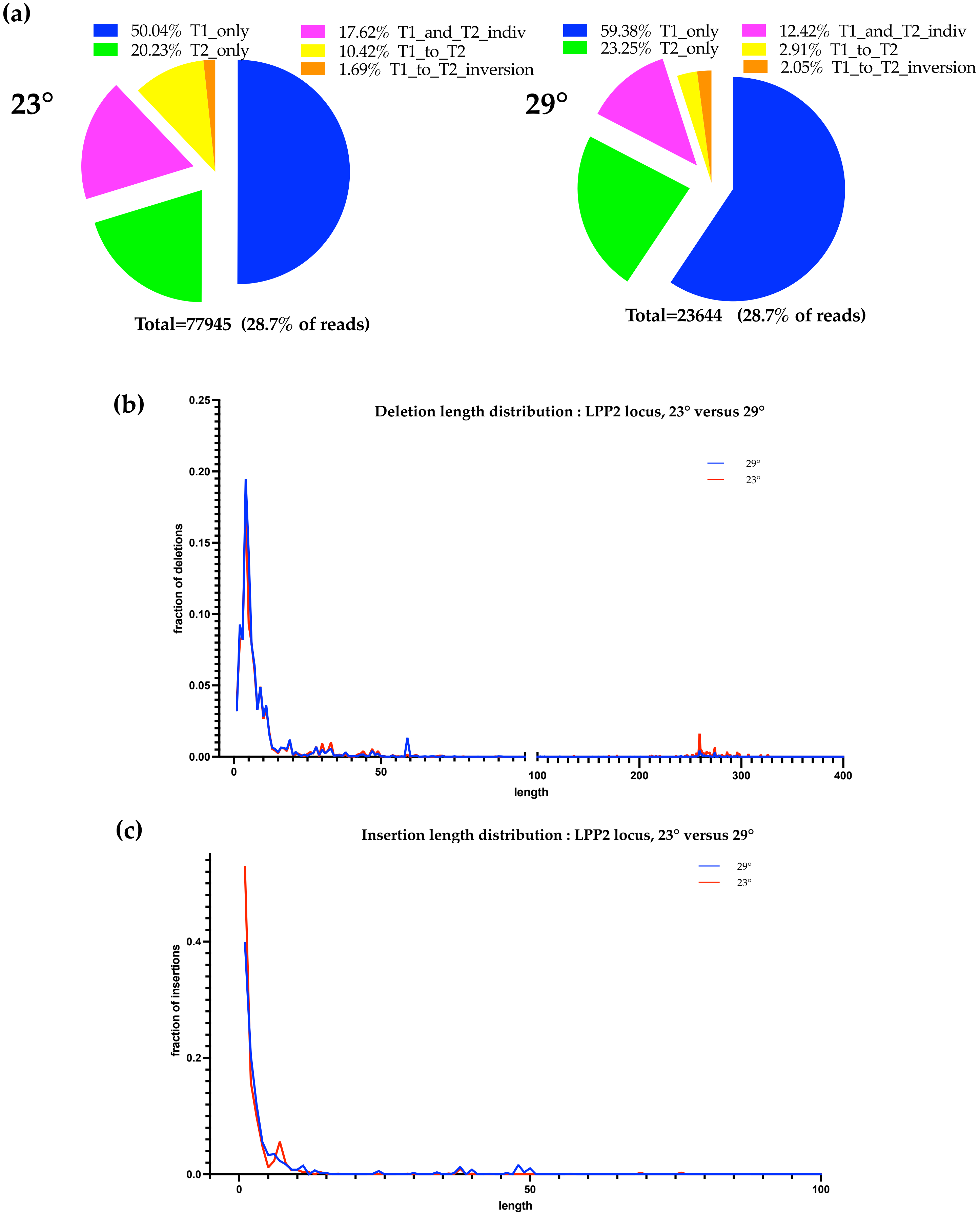
(a)
50.04% T1_only
20.23% T2_only
17.62% T1_and_T2_indiv
10.42% T1_to_T2
1.69% T1_to_T2_inversion
23°
Total=77945 (28.7% of reads)
59.38% T1_only
23.25% T2_only
12.42% T1_and_T2_indiv
2.91% T1_to_T2
2.05% T1_to_T2_inversion
29°
Total=23644 (28.7% of reads)
(b)
Deletion length distribution : LPP2 locus, 23° versus 29°
29°
23°
fraction of deletions
0.25
0.20
0.15
0.10
0.05
0.00
0
50
100
200
300
400
length
(c)
Insertion length distribution : LPP2 locus, 23° versus 29°
29°
23°
fraction of insertions
0.4
0.2
0.0
0
50
100
length

### Figure S4

(a)

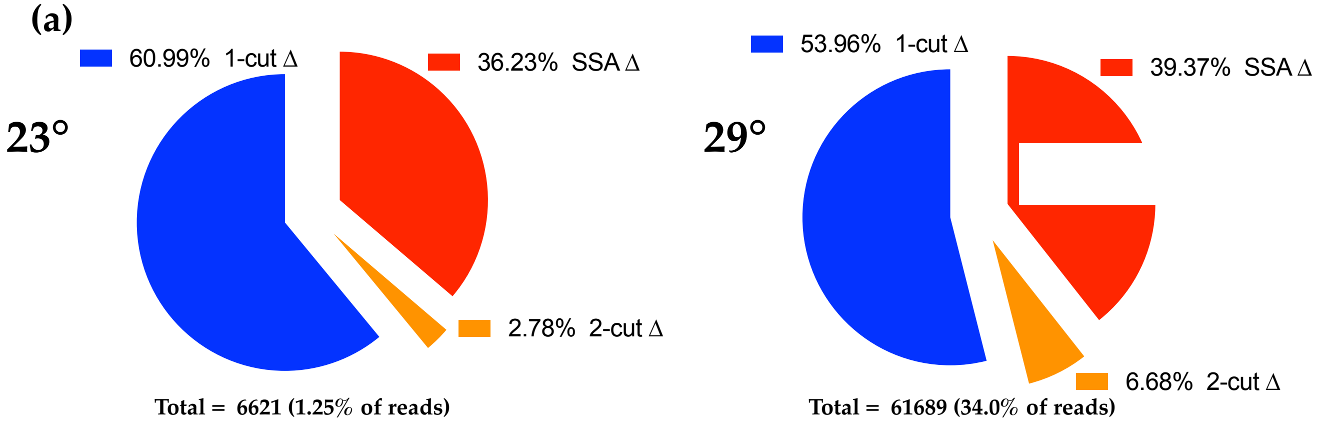


(b)

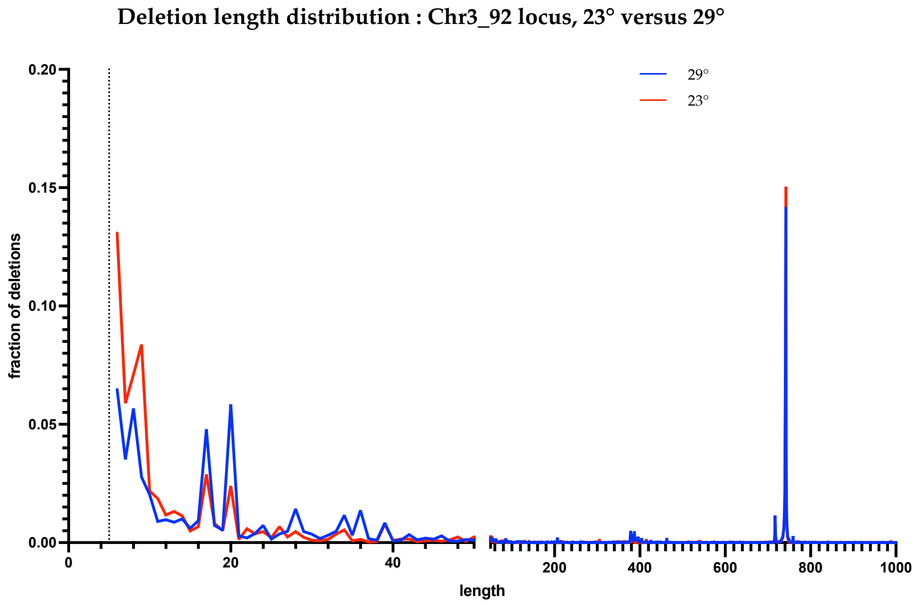


(c)

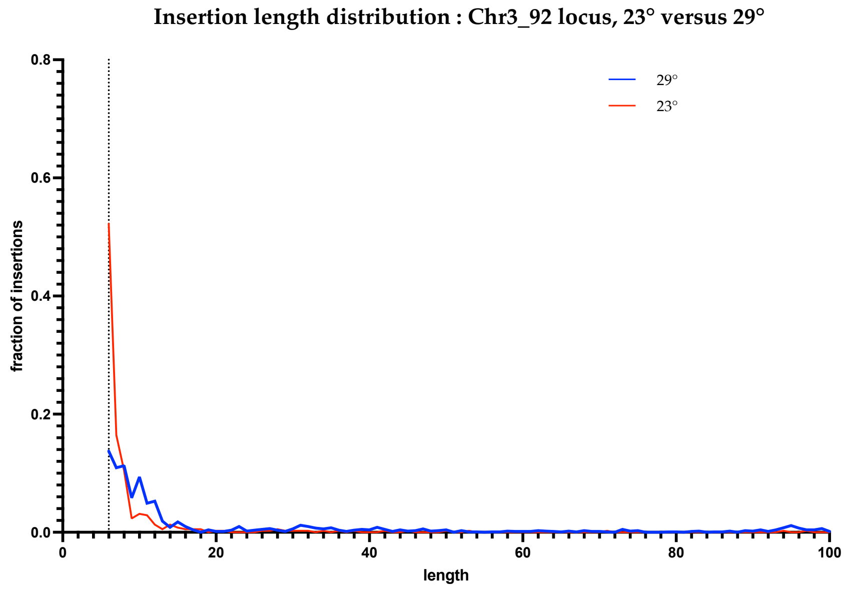

### Supporting Tables

| **Table S1: DNA sequences used for sgRNA constructs**<br>crRNA sequences are in upper-case, fnCas12a direct repeats are magenta, BbsI restriction sites are red | |
|---|---|
| gRNA_LPP2_top | 5'-agatAGCTCGACCAAGTTCCCTTTGTTaatttctactgttgtagatGTTGGAGAAGATATGCTCACTGA-3' |
| gRNA_LPP2_bottom | 5'-aaaaTCAGTGAGCATATCTTCTCCAACatctacaacagtagaaattAACAAAGGGAACTTGGTCGAGCT-3' |
| gRNA_Chr3_92_top | 5'-agatATAGATGCACGAACACATCAACAaatttctactgttgtagatTAGTATATGAACTAGTTTCACGG-3' |
| gRNA_Chr3_92_bottom | 5'-aaaaCCGTGAAACTAGTTCATATACTAatctacaacagtagaaattTGTTGATGTGTTCGTGCATCTAT-3' |

| **Table S2: List of primers used in this study** | |
|---|---|
| LPP2_MiSeq_P5tag_For | 5'-TCGTCGGCAGCGTCAGATGTGTATAAGAGACAGGATTCTTGTTTGTGGCCATTGTGCA-3' |
| LPP2_MiSeq_P7tag_Rev | 5'-GTCTCGTGGGCTCGGAGATGTGTATAAGAGACAGCGGGACAGCCCAGAAAGGA-3' |
| Chr3_92_MiSeq_P5tag_For | 5'-TCGTCGGCAGCGTCAGATGTGTATAAGAGACAGTTGTTGGATATAGTATGATGTTGGAC-3' |
| Chr3_92_MiSeq_P7tag_Rev | 5'-GTCTCGTGGGCTCGGAGATGTGTATAAGAGACAGAACAGAAAATTTGATTATTGGAGGG-3' |
| LLP2_Nanopore_For | 5'-GAGAAGGTTTCGTGGGGAGC-3' |
| LLP2_Nanopore_Rev | 5'-ACGCTCTCTAGCTCTGTTTG-3' |
| Chr3_92_Nanopore_For | 5'-ATTCGATTGTACGTTGTGGCTG-3' |
| Chr3_92_Nanopore_Rev | 5'-CCTCTGTTCGTTGATTTTTGGGAAC-3' |
